# Corticosterone as a potential confounding factor in delineating mechanisms underlying ketamine’s rapid antidepressant actions

**DOI:** 10.1101/2019.12.30.891333

**Authors:** Lauren Wegman-Points, Brock Pope, Allison Zobel, Lori Semke, Eric Wauson, Vanja Duric, Li-Lian Yuan

## Abstract

Although ketamine represents a new line of antidepressants with unique clinical advantages, its use as a long-term treatment has limitations, particularly its dissociative/psychotomimetic effects and abuse potential. In rats, we observed that a subanesthetic dose of ketamine (10mg/kg) induced a 3-fold increase in corticosterone (CORT) levels in both serum and brain tissue, within an hour of administration. This increase took place in both male and female rats, in both naïve and stressed animals. However, no CORT increase was detected in rats injected with (2R, 6R)-hydroxynorketamine (HNK), an active metabolite of ketamine, that is believed to contribute to ketamine’s antidepressant effect. In response to the release in CORT, ketamine injected animals displayed a significant increase in the expression of *sgk1*, a downstream effector of glucocorticoid receptor signaling, in the hippocampus indicating the initiation of a transcriptional program. We hypothesized this surge in CORT release was a manifestation of stress experienced by the rat in response to ketamine’s psychotropic effects. When sensory perception was blocked under isoflurane anesthesia, administration of ketamine did not increase circulating CORT levels as compared to animals injected with saline. Taken together, ketamine administration triggers a behavioral stress response that has downstream molecular consequences. The resulting CORT release, virtually concurrent with the timing of ketamine’s rapid-acting antidepressant actions, necessitates the consideration of this pathway’s potential involvement when trying to dissect out the relevant molecular mechanisms underlying ketamine’s action.

## 1. Introduction

Ketamine, synthesized as a phenylcyclidine (PCP) derivative over 50 years ago, has long been used as general anesthetic acting primarily through blockade of NMDA receptors in the brain[1]. However, ketamine, at sub-anesthetic doses, has recently emerged as a clinical rapid acting antidepressant (RAAD). Accumulating evidence supports ketamine’s efficacy in the treatment of major depressive disorder (MDD) and other anxiety disorders with the administration of subanesthetic doses of ketamine[2-5]. Ketamine not only offers quick relief from depressive symptoms and suicidal ideation within a few hours, it is also effective in a large subset of the treatment resistant population who do not respond to current mainstream monoamine therapies [6]. Despite ketamine’s unique clinical advantages, its use as a safe, long-term treatment has limitations. Even at subanesthetic doses, ketamine produces psychotomimetic and dissociative effects. These effects increase the potential for abuse, largely restricting administration to a clinical setting. The majority of the studies examining the effects of long-term repeated dosing of ketamine have been on chronic abusers, thus the specific effects of long-term low dose use for treatment of TRD are largely unknown.

The adverse effects associated with ketamine are not only a clinical concern, but also a critical barrier to elucidating the molecular mechanisms underlying its antidepressant actions. Animal model studies have identified multiple mechanisms related to ketamine’s rapid action, including rapid enhancement of glutamate transmission and production of BDNF neurotrophic factors, activation of mTOR signaling pathways in the prefrontal cortex (PFC), suppression of neuronal firing in lateral habenula, and potential contributions of a ketamine metabolite hydroxynorketamine (HNK)[7-10]. Together, these molecular and cellular events are proposed to lead to rapid synaptogenesis and reversal of neural atrophy. However, consensus in neural mechanisms has yet to be developed and major questions remain, such as whether ketamine’s antidepressant action is NMDAR dependent or independent[6, 11]. Further complicating matters, ketamine treatment also leads to a host of biochemical alterations including increased glucocorticoid circulation and hormonal responses (e.g. prolactin) that occur in addition to ketamine’s NMDAR antagonism [12-14]. The transient increase in circulating glucocorticoids in response to the antidepressant dose of ketamine peaks between 30 to 120 minutes of treatment onset, depending upon the route of administration, and returns to baseline within a couple hours of treatment completion, in both rodent models and human subjects [12, 13, 15].

Glucocorticoids mediate the body’s response to stressful situations. Cortisol, corticosterone (CORT) in rodents, is released from the adrenal cortex in response to pathways initiated in the hypothalamus and pituitary gland [16]. Research has shown that depression and the hypothalamic-pituitary-adrenal (HPA) axis are intimately intertwined [17]. In fact, several studies document an increase in volume in both the pituitary and adrenal glands in patients with MDD [16]. CORT signaling in acute bouts, has physiological properties and contributes to learning and coping skills [18]. However, when the signaling persists, due to prolonged stress or when coping is not accomplished, the effects of CORT become pathological[19].

We feel both the role of glucocorticoids in MDD and the timing of this release warrant further research into this effect of ketamine administration. The timing of this glucocorticoid release is completely concurrent with the initiation of ketamine’s therapeutic effects which occur within two hours of administration [13]. Focusing within that timeframe, here we examine the effect in both male and female rodents, downstream signaling pathways, and begin to examine the mechanism of its generation. We also discuss whether this phenomenon could contribute to the RAAD effects of ketamine, or if it merely exists as a confounding factor.

## 2. Materials and Methods

### 2.1. Animals

8-10 week old Sprague-Dawley rats male and female were obtained from Charles River (Wilmington, MA). Male and female rat cohorts were done separately. The animals were pair-housed under a 12-h light/dark cycle (lights on at 6:00am) with controlled temperatures and humidity. Food and water were administered ad libitum. Body weight was recorded on a weekly basis, and overnight fluid consumption was monitored twice per week. The use of animals for these studies was approved by the Des Moines University Institutional Animal Care and Use Committee.

### 2.2. Drugs

(2R,6R)-hydroxynorketamine hydrochloride (R&D Systems, Minneapolis, MN) was dissolved in saline at 10mg/ml immediately prior to use. Prior to administration, Ketamine hydrochloride (Ketaset, Zoetis, US) was diluted from 100mg/ml to 10mg/ml in saline. Corticosterone hemisuccinate (4-pregen-11B 21-DIOL-3 20-DIONE 21 hemisuccinate) (Steraloids, Newport RI) was added to tap water at concentration of 64.4mg/L (producing a final concentration of 50ug/ml CORT), brought to pH 12.5 using NaOH, and allowed to dissolve overnight at 4°C on a stir plate. The pH was adjusted to between 7.2-7.5 with HCl before administration to the animals. 25ug/ml and 12ug/ml doses were prepared in the same manner with 32.2mg/L and 16.1mg/L of corticosterone hemisuccinate respectively.

### 2.3. Chronic stress model

CORT was administered to the rats through their drinking water according to Gourley et al. [22]. The rats were provided with water containing 50ug/ml of CORT for 2 weeks. A freshly prepared solution was provided every 3 days. Daily dosages of were calculated using overnight water consumption values and body weight. 5-7mg/kg was considered the minimum effective dosage. After the initial 2 weeks, the dosage was tapered off with 3 days to 25ug/ml followed by 3 more days at 12.5 ug/ml. The rats were returned to regular drinking water for 3 days to clear the system of exogenous CORT before injection and sacrifice.

### 2.4. Blood and tissue collection

Animals were euthanized by decapitation 1 hour after receiving an intraperitoneal (i.p.) injection of saline, ketamine or HNK. Treatment groups were staggered throughout the day to avoid natural oscillations in circadian pathways from affecting any one treatment group. Trunk blood was collected in serum collection tubes (BD Bioscience) and allowed to clot 15-30 min before centrifugation. Brain tissue was dissected, snap frozen using liquid nitrogen, and stored at -80°C until use.

### 2.5. Blood collections under sedation

8-12 week old animals were sedated using 3% isoflurane in a ventilated chamber. After initial sedation, the animals were maintained under anesthesia at 1.5% isoflurane through a nosecone. A heating pad was used to maintain body temperature. Hind limbs were shaved and treated with a chemical depilatory to allow visualization of the saphenous vein. A 23.5-gauge needle was used to puncture the saphenous vein and blood was collected using a serum capillary blood collection tube (Microvette 200, Sarsedt, Germany). After the initial blood collection, an i.p. injection of saline or ketamine (10mg/kg) was administered. The animals remained under anesthesia for 60 minutes after the injection when blood was collected again through the saphenous vein. Animals were allowed to recover and returned to their cages.

### 2.6. Measurement of endogenous corticosterone levels

For measurement of endogenous corticosterone (CORT) levels in serum and cortical brain tissue, we utilized the DetectX CORT ELISA kit (Arbor Assays, Ann Arbor, MI). For the cortical tissue, a separate steroid extraction procedure was required before CORT measurement. Briefly, approximately 50mg of cortical tissue was manually homogenized using a pellet pestle and incubated in ethyl acetate (1ml/0.1g tissue) with agitation for 30 min. Samples were spun at 5,000rpm for 15 min at 4°C and the supernatant was collected. The supernatant was evaporated under a vacuum (without heat) using a SpeedVac for 45 minutes and stored in a dessicator at -20°Cuntil use. Samples were reconstituted in 100% EtOH immediately prior to analysis. A kit-provided dissociation buffer was used to extract CORT in the serum samples. The assay was performed according to the manufacturer’s instructions, which each sample run in duplicate. Samples were read at 450nm and compared to a standard curve using a 4 parameter logistical regression model.

### 2.7. RNA extraction and qPCR

Total RNA was extracted from the entire hippocampus using the RNeasy Lipid Tissue Mini kit (Qiagen, Germantown, MD). cDNA was generated using the High Capacity cDNA Reverse Transcription Kit (Applied Biosystems, Waltham, MA) using random hexamer primers. qRT-PCR was performed on a CFX Connect Real Time PCR Detection System (Bio-Rad, Hercules, CA) utilizing iTaq Universal SYBR Green Supermix (Bio-Rad, Hercules, CA). B-Actin and GAPDH were both used as control genes in all experiments. B-Actin: (PrimeTime® IDT, Coralville, IA) Forward 5’-TCACTATCGGCAATGAGCG-3’, Reverse 5’GGCATAGAGGTCTTTACGGATG-3’. Primer 3 software was utilized to design SGK-1 and GAPDH primer sets. SGK-1 Forward 5’-GTGCCTTGCTAGAAGAACCTTTCC-3’, Reverse 5’-CTCACCTCCTCCAAGTCCCTCTC-3’. GAPDH Forward 5’-GTCATGAGCCCTTCCACGATGC-3’, Reverse 5’-ACAACTCCCTCAAGATTGTCAGC-3’[20, 21].

### 2.8. Statistics

One-way analysis of variance (ANOVA) was used to analyze the concentrations of CORT between control and stressed treatment groups with males and females analyzed separately. The 1-hour timepoint qPCR results were also analyzed by a one-way ANOVA using the ΔCt values. An unpaired student’s t-test was utilized in comparison of CORT levels in all naïve and 24-hour animal experiments. Graphpad Prism was used to perform all statistical analyses.

## 3. Results

### 3.1. Increase in circulating CORT with ketamine treatment

To learn more about the ketamine’s initial transient molecular pathways, we utilized a well-established oral CORT stress model to sensitize the system and compare the molecular effects of treatment with ketamine versus its metabolite HNK [22]. Recent work provided evidence that HNK on its own is sufficient for the RAAD effect in rodents, although subsequent work has challenged this assertion[7, 23, 24]. HNK does not induce psychotropic side effects, making it an attractive RAAD candidate for further investigation [11]. CORT (50μg/ml) was administered to experimental rats through their drinking water to achieve an effective working dosage of 5-7mg/kg. Control animals were supplied with normal drinking water. Male and female animals were examined separately in independent cohorts. One hour prior to sacrifice, stressed animals were injected intraperitoneally (i.p.) with either saline, ketamine (10mg/kg) or HNK (10mg/kg) while control animals received saline. ELISA analysis of trunk blood collected at sacrifice, indicated a substantial surge in CORT release only in the animals treated with ketamine (Figure 1A). Ketamine-injected male rats showed several folds more circulating CORT (532.8 ng/ml ± 47.3; F(3, 39)=19.38, p<0.0001) than water control animals injected with saline (136.1 ng/ml ± 35.6)(Figure 1A). Neither of the remaining stressed groups, saline (189.4 ng/ml ± 51.6) or HNK (108.7 ng/ml ± 33.3) displayed a significant increase in circulating CORT (Figure 1A).

**Figure 1.**
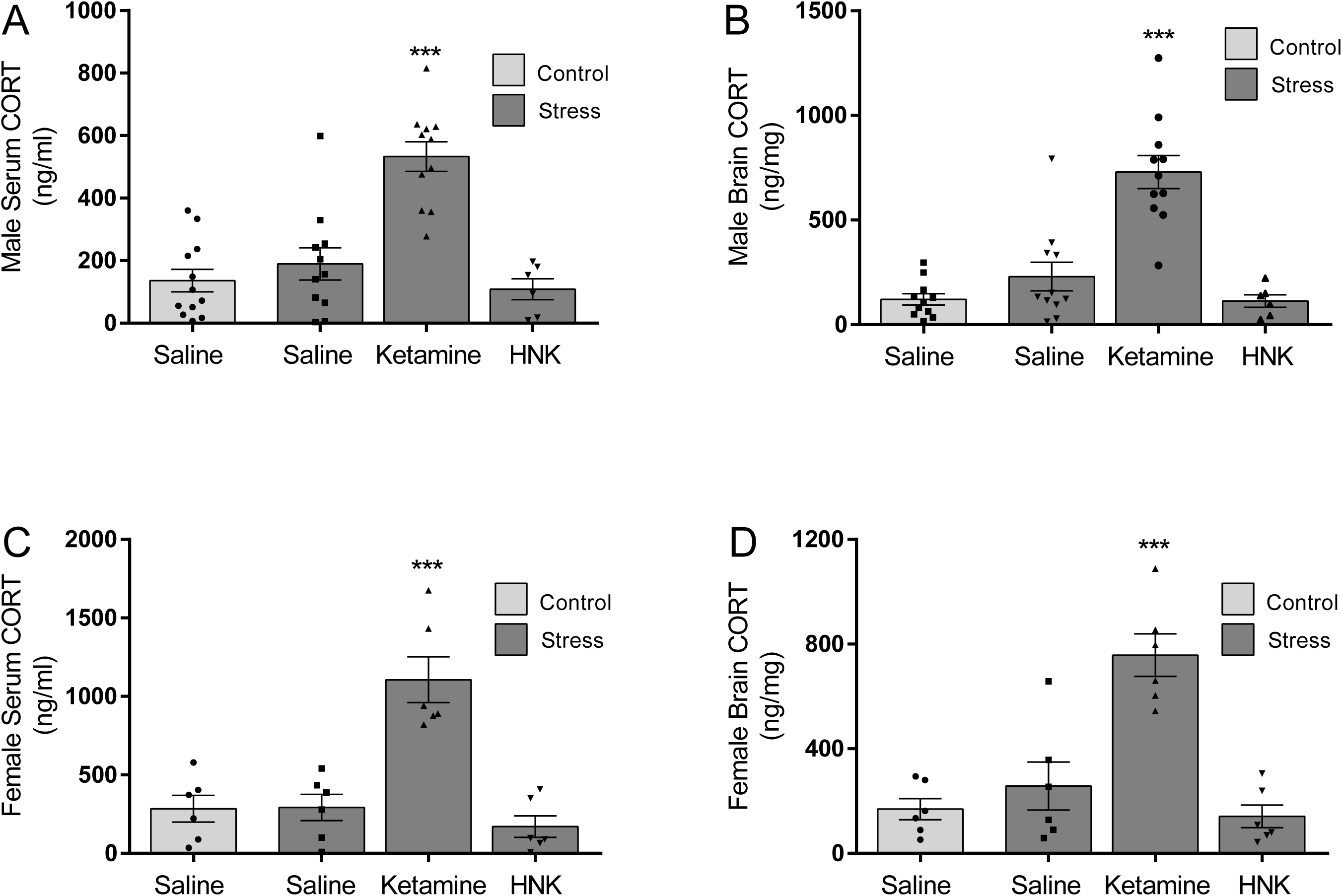
Measurement of CORT levels in serum and brain tissue one hour post drug treatment. Rats of both sexes exposed to chronic stress received injection of saline, ketamine (10 mg/kg) or HNK (10 mg/kg). Control rats received only saline. **A**. Circulating CORT levels in male rats. **B**. Cortical brain levels of CORT in male rats. **C**. Circulating CORT levels in female rats. **D**. Cortical brain levels of CORT in female rats. (Control n=11, Stress/Saline n=11, Stress/Ketamine n=11, Stress/HNK n=6; ***p<0.0001, one-way ANOVA).

Females, while displaying higher overall levels of CORT, exhibited the same pattern as the males in ketamine-treated animals (1106 ng/ml ± 146.0; F(3, 20)=18.70, p<0.0001) with levels almost 4 times higher than control animals (284.2 ng/ml ± 84.3) and the other stressed groups, saline (292.2 ng/ml ± 83.2) and HNK (170.3 ng/ml ± 68.2)(Figure 1C) These surges in CORT seen in both male and female rats were well above the daily rise in CORT levels seen due to normal circadian oscillations.

### 3.2. CORT increase in brain tissue

Next we examined CORT levels in cortical brain tissue collected at the 1 hour post-injection timepoint. We performed a steroid extraction on the tissue and analyzed the extracts by ELISA. The upsurge in circulating CORT translated to a concurrent increase in CORT levels in the brain of the ketamine-treated animals. Male ketamine-treated rats showed drastically higher CORT levels (729.9 ng/mg ± 78.75; F(3, 35)=23.64, p<0.0001) versus controls (120.9 ng/mg ± 26.74) and the stressed treatment groups, saline (229.6 ng/mg ± 68.12) and HNK (113.5 ng/mg ± 30.0) (Figure 1B). Female ketamine-treated animals also displayed an increased level of CORT in the brain relative to the other treatment groups. Interestingly, unlike in the circulating CORT, in cortical brain tissue, the female treatment groups showed levels more in line with their male counterparts. Female ketamine-treated animals brain CORT levels (758.0 ng/mg ± 81.46; F(3, 20)=17.90, p<0.0001) were still significantly higher levels than controls (169.2 ng/mg ± 40.57) and the stressed treatment groups, saline (257.8 ng/mg ± 92.0) and HNK (141.2 ng/mg ± 43.2) (Figure 1D).

### 3.3. Increase in circulating CORT in naïve animals

In order to determine if our stress protocol had any effect on the animal’s CORT response to ketamine treatment, we performed a similar set of experiments on naïve animals. Animals were given an i.p. injection of saline or ketamine and euthanized at 1-hour to collect blood and tissue. Circulating CORT levels were significantly elevated in the animals treated with ketamine versus those receiving saline. These results indicate that our previous results were not an artifact of our stress protocol.

### 3.4. Analysis of CORT regulated transcripts

To investigate the molecular consequences of the ketamine-induced increase in brain CORT levels, we used qRT-PCR to look for transcriptional changes in the glucocorticoid receptor downstream effector serum glucocorticoid kinase 1 (*sgk-1)*. We chose the hippocampus for our analysis, because as part of the limbic system, it is integrally involved in the animal’s stress response. *Sgk-1*, is a serine/threonine kinase implicated in the cellular stress response and neuronal excitability [25]. Male ketamine injected animals displayed a significant decrease in ΔCt values (F(3, 20) = 4.4, p<0.015) versus controls and stressed treatment groups. This translated to an almost 2-fold increase (1.88 ± 0.16) in *sgk1* transcript over controls in response to ketamine administration, versus stressed animals injected with saline (1.02 ± .25) or HNK (1.40 ± .27) (Figure 3A). Females also displayed a significant decrease in ΔCt values (F(3, 20) = 13.02, p=0.0001) in ketamine-treated animals versus controls and other treatment groups. This represented a 1.5-fold increase (1.51 ± .19) in *sgk-1* expression over saline water controls, a 1.8-fold increase over stressed saline (0.82 ± .11) and 2.4-fold over stressed HNK (0.62 ± .04) groups (Figure 3B). We also examined *sgk-1* mRNA levels in stressed male rats at 24 hours to see if the levels had returned to baseline. 24 hours after injection with either saline or ketamine (10mg/kg) *sgk-1* mRNA levels were indistinguishable between groups (Figure 3C). Both male and female 1-hour treatment groups and the male 24-hour treatment groups were also examined for differences in FKBP5 expression, another glucocorticoid receptor downstream target, however there was no change in expression at either the 1-hour or 24-hour timepoints for any of the groups (data not shown). This was likely due to the timing of the tissue collection, and additional timepoints would allow for a more thorough characterization of CORT’s downstream effects.

**Figure 2.**
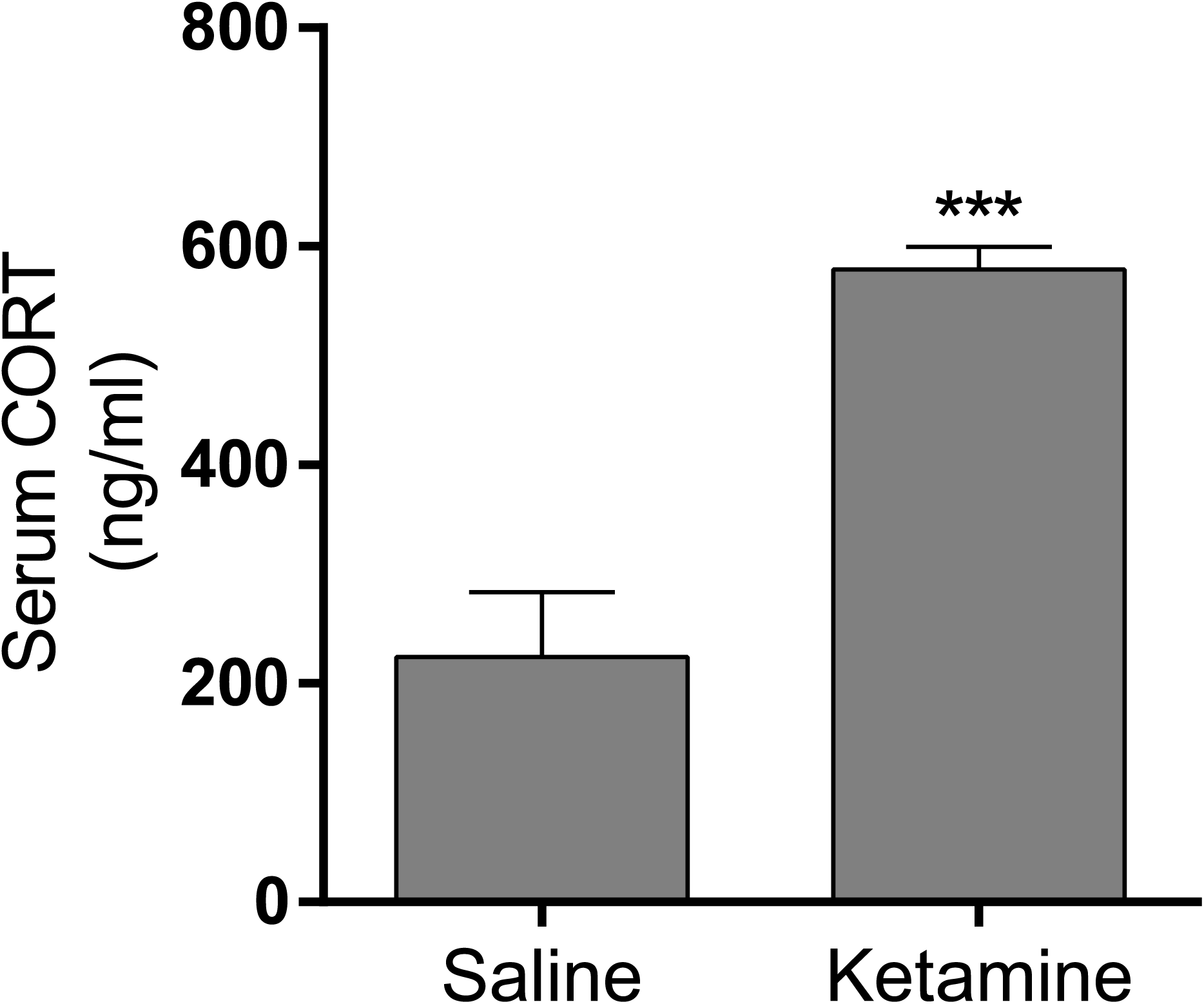
Measurement of serum CORT levels one hour post drug treatment of naïve rats (saline n=5, ketamine n=5). Ketamine-induced surge in CORT release was also observed in naïve rats. (***p<0.0005, one-way ANOVA)

**Figure 3.**
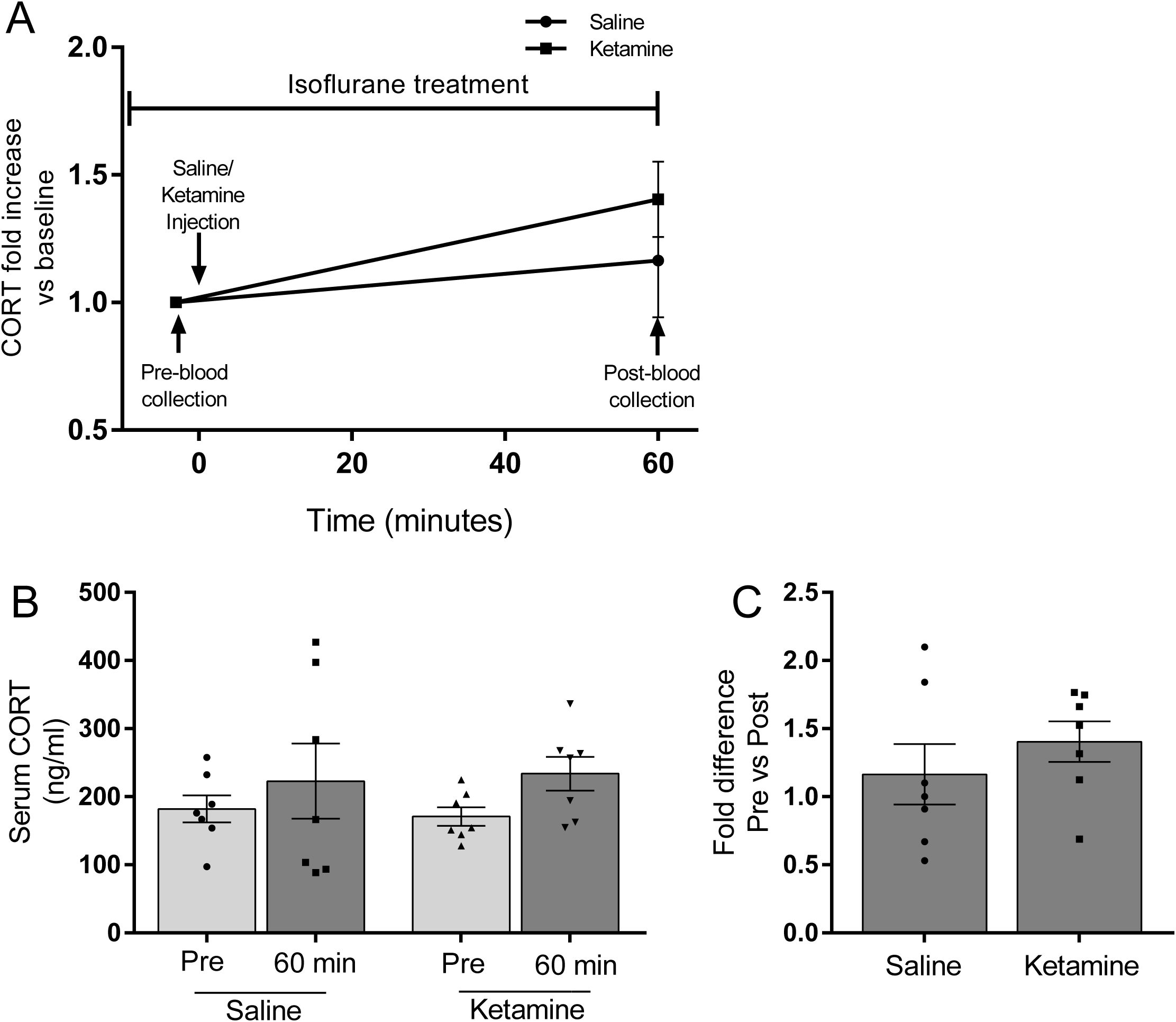
Expression of <u>S</u>erum/<u>G</u>lucocorticoid Regulated <u>K</u>inase (*sgk1)* in the hippocampus (HPC) of male and female rats at 1 hour and 24 hours post injection. Rats of both sexes exposed to chronic stress. Sgk1 levels increase in response to the CORT release and return to baseline at 24 hours. **A**) *Sgk1* expression in HPC of male rats 1 hour after injection (n=6 for each condition). **B**) *Sgk1* expression in HPC of female rats 1 hour after injection (n=6 for each condition). **C**) *Sgk1* expression in HPC of male rats 24 hour after injection (n=4 for each condition). All values are presented as a fold change from stress control animals injected with saline. (** p<0.01 ANOVA on ΔCt values).

### 3.4. Ketamine effects on circulating CORT under anesthesia

Within minutes of administration of 10mg/kg ketamine, freely mobile conscious rats begin to exhibit locomotor deficits, including an unsteady gait and falling over onto their side. We hypothesized this represents an outward manifestation of the dissociative effects described by human subjects. We reasoned that this novel experience could be generating anxiety and confusion in the animal and may be responsible for the surge in CORT we see. To determine whether the CORT surge is in response to the animal’s conscious perception of the psychotropic effects of ketamine, we examined the effects of ketamine on CORT under anesthesia. We anesthetized naïve male rats with isoflurane. Our experimental design is depicted in Figure 4A. While the animals were under anesthesia, we took blood at baseline, then injected via i.p. either saline or ketamine (10 mg/kg). After 1 hour, we took another sample of blood to examine CORT levels in response to the saline or ketamine injections. There was no statistical difference between the saline and ketamine groups in pre-injection CORT levels (181.9 ± 19.8 vs 170.9 ± 13.45, p<0.6541), or in the post-injection CORT levels (222.9 ± 55.2 vs 233.8 ± 24.9, p<0.8595) (Figure 4B). There was a modest upwards trend in the ketamine pre versus post CORT (Mean of Diff 62.87 ± 26.97, p<0.056) levels, however this trend did not reach significance, and the fold changes seen in the ketamine group did not approach the levels seen in conscious animals (Figure 4C).

**Figure 4.**
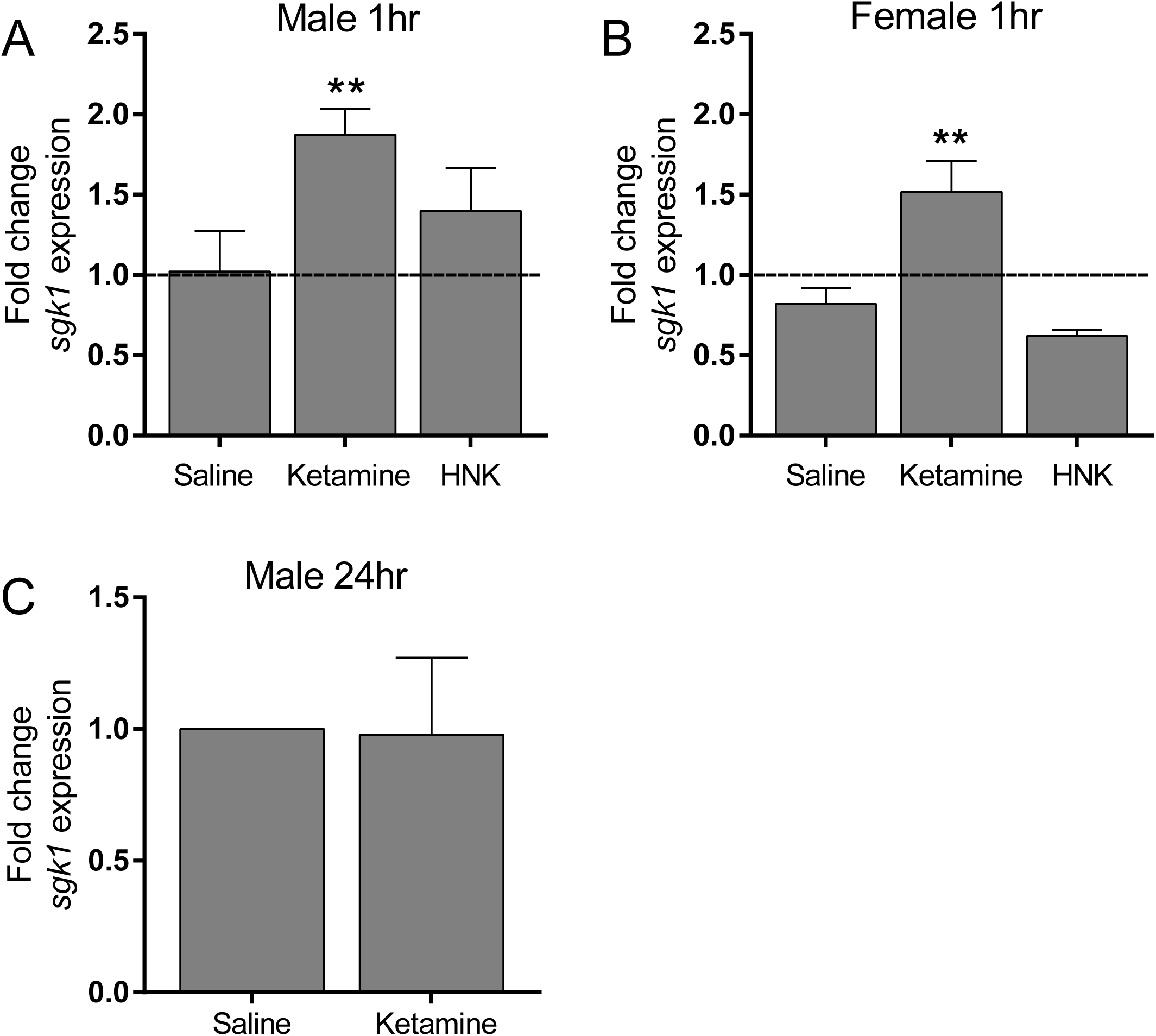
Circulating CORT levels of male rats under isoflurane anesthesia before and after drug treatment. Removing conscious perception of dissociative effects appears to mitigate the ketamine-mediated CORT release. **A**. Timeline showing the experimental design. **B**. There are no significant differences between pre-injection and post-injection CORT levels for saline or ketamine (n=7 for both groups). **C**. Fold differences between pre and post CORT levels after injection.

## 4. Discussion

Here we have further characterized a surge in glucocorticoid release upon administration of a subanesthetic dose of ketamine. Although previously reported, we felt this phenomenon warranted further examination both into its effects and the mechanism behind its generation. This increase in CORT is seen across both sexes, and is specific to ketamine treatment, as we did not see an increase with the injection of saline or HNK. The effect is present whether looking at naïve animals, or those within a stress paradigm. We also show the downstream effects on transcription in the hippocampus, a region shown to be highly relevant in the manifestation of MDD. Finally, we begin to offer a potential mechanism for this increase in circulating CORT, namely as a behavioral response to the experience of ketamine’s well-documented psychotomimetic and dissociative effects.

Because dysregulation of glucocorticoid signaling in MDD is well-cited in the literature, the initial impetus for examining the circulating levels of CORT, was to confirm normal activity of the HPA axis following our stress protocol and document any differences that could affect our future experimental results[26]. Not aware of previous publications documenting increased glucocorticoid levels after ketamine administration, we were surprised to see the substantial rise in circulating CORT levels specific to the administration of ketamine. Of particular interest, was that the same effect was not seen with HNK administration. Our initial experiments were done in male rats, so we went on to confirm the same effect in females. Females displayed an overall higher level of CORT, which agrees with previous publications, but the ratios of CORT levels between ketamine and the other treatment groups were remarkably similar[27]. In both sexes, ketamine increased circulating CORT levels almost 4-fold when compared to untreated saline-injected controls.

Glucocorticoids must pass through the blood brain barrier to exert their negative feedback effects on the HPA axis, so we were not surprised to confirm a similar increase in CORT in cerebral cortical tissue collected from the same animals [28]. One major pathway CORT utilizes to exert its downstream effects is through activation of the glucocorticoid receptor [17]. Upon CORT binding, the glucocorticoid receptor complex exerts transcriptional effects on genes with promoters containing a Glucocorticoid Response Element (GRE) [28]. By analyzing levels of *sgk1* mRNA, a GRE containing gene, in hippocampal tissue from our animals, we were able to confirm both the presence and activity of increased CORT levels in the brain. This is particularly relevant to RAAD research as the hippocampus, part of the limbic system and the primary stress response circuit, has been strongly implicated in the pathophysiology of MDD [26] Clinical and postmortem studies of MDD patients show evidence of atrophy, reduced neurogenesis and an overall reduction in hippocampal volume [18, 29]. Ketamine administration initiating glucocorticoid transcriptional pathways in key MDD nuclei could possibly affect research efforts into other pathways pertinent to the RAAD effects and should be considered when analyzing results.

How does a canonical NMDAR antagonist exert effects on CORT release? There are potential direct biochemical mechanisms that could generate the increase in CORT levels seen in our ketamine-treated animals. Sympathomimetic properties acting on the adrenal gland or direct binding to the pool of NMDA-Rs in the hypothalamus [12]. However, we assert that it is not a direct biochemical effect of ketamine binding. We suggest it is part of an acute behavioral stress response the experience of the dissociative effects of ketamine. Upon administration of ketamine, but not of HNK, the animal’s motor coordination begins to become dysregulated. We assert this is a physical representation of the dissociative and psychotomimetic effects experienced with ketamine administration. Our data suggests this novel and potentially stressful experience causes an acute CORT response. Removing the animal’s experience of these effects through anesthesia prevented the ketamine-induced increase in circulating CORT. While we did see a slight non-significant rise in CORT levels, this was likely due to either the isoflurane or repeated blood draw and did not approach the levels seen in conscious animals. Our choice of isoflurane as our anesthetic was based on research showing it to have minimal effect on circulating CORT levels in male rats [27, 30]. With this choice, we hoped to mitigate the effects of anesthesia on our results, however we cannot completely rule out the possibility that the animal’s reduced CORT response is an effect of the isoflurane on the HPA axis.

The concurrent increase in CORT with ketamine administration raises the question as to whether the activation of the HPA axis and its downstream pathways, are a necessary component to its antidepressant effect. While warranting additional investigation, several human studies have shown a link between the severity of the dissociative effects and the strength of the antidepressant effect [5, 31, 32]. Interestingly, the antidepressant effect is not seen with anesthetic doses of ketamine in mice[8]. HNK does not cause dissociation, nor does it increase circulating CORT levels, yet Zanos et al still reported rapid antidepressant effects using this compound. However, as stated previously, this assertion that HNK is necessary and sufficient to generate RAAD effects has come under scrutiny [24]. Future experiments to see if animals treated with subanesthetic doses of ketamine under anesthesia experience a sustained antidepressant effect at 24 hours might help answer the question of whether dissociation is relevant to ketamine’s AD effect.

Uncovering the salient pathway(s) leading to ketamine’s sustained antidepressant effects could possibly lead to the generation of treatments with the same RAAD effects and avoiding the undesirable side effects. However, there remains the possibility that the sustained antidepressant effects cannot be uncoupled from the dissociative effects. Here we have shown that outside of ketamine’s documented effects on glutamate signaling, there is a secondary behavioral component that has molecular consequences. The conscious perception of ketamine’s dissociative effects appears to initiate its own set of secondary effects adjacent to the NMDA-R antagonist pathway.

Could these pathways converge, ultimately working in synchrony to increase synaptic strengthening and the sustained relief of depressive symptoms? At the very least, because CORT signaling is implicated in so many pathways relevant to a depressive phenotype, the surge in CORT should be considered when analyzing any results from ketamine administration. With such a thrust in ketamine research, and no definitive answers at hand, it is becoming increasingly apparent that we must consider that ketamine’s antidepressant effect may lie in a distinct combination of events instead of a single receptor target.

## Declaration of Interest

The authors declare no competing financial interests.

## Acknowledgements

We thank Kiefer Kious and Dr. Noah Marcus for generously providing access to and training for using their ventilator machine, and Dr. Chunfa Jie for help with statistical analysis. This work was supported by National Institutes of Health grants (MH115396 and MH108043 to L.Y.).

